# A model-based analysis reveals three-dimensional genome organization heterogeneity and functional sub-compartments in cell populations

**DOI:** 10.1101/2022.06.27.497761

**Authors:** Xiaobin Zheng, Joseph R. Tran, Yixian Zheng

## Abstract

The computational inference of genome organization based on Hi-C sequencing has greatly aided the understanding of chromatin and nuclear organization in three dimensions (3D). However, existing computational methods used to infer sub-compartments from Hi-C data fail to address the cell population heterogeneity. Here we describe a model-based method, called CscoreTool-M, which uses probabilistic modeling to build multiple 3D genome sub-compartments from Hi-C data. The compartment scores inferred using CscoreTool-M directly represents the probability of a genomic region locating in a specific sub-compartment. Compared to published methods, CscoreTool-M is more accurate in inferring local sub-compartment containing heterochromatin marked by Histone lysine trimethylation (H3K27me3) surrounded by the actively transcribed euchromatic regions. The compartment scores calculated by CscoreTool-M also help to quantify the levels of heterogeneity in sub-compartment localization within cell populations for different genomic regions. By comparing proliferating cells and terminally differentiated non-proliferating cells, we show that the proliferating cells have higher genome organization heterogeneity, which is likely caused by cells at different cell-cycle stages. By analyzing 10 sub-compartments, we found a sub-compartment containing chromatin potentially related to the early-G1 chromatin regions proximal to the nuclear lamina in HCT116 cells, suggesting the method can deconvolve cell cycle stage-specific genome organization among asynchronously dividing cells. Finally, we show that CscoreTool-M can further identify sub-compartments that contain genes enriched in housekeeping or cell-type-specific functions.

## Introduction

One important question in biology pertains to how DNA is packaged and organized in the small nuclear space of eukaryotes so that gene expression can be temporally and spatially regulated during organism development and homeostasis. The nucleus is a complicated structure with many functional subunits, such as the nuclear speckles, the nucleolus, and the nuclear lamina. These units have been recognized as organization hubs of the genome for specific functions. How these hubs are organized with respect to one another and to the 3D chromatin interactions remains, however, poorly understood. As a result, it is unclear how the different genomic units work together to regulate gene expression in the context of organism development and function.

The genomic sequences associated with the nuclear lamina, speckles, and nucleoli have been mapped (1–3). The genes in the nuclear lamina-associated chromatin domains (LADs) are largely silenced and the LADs are generally gene-poor with overall repressive epigenetic features. Nuclear speckles are enriched for transcribed genes that create a transcriptionally active environment. Thus, chromatin associated with the speckles show active epigenetic features. It remains unclear if the actively transcribed house-keeping genes and cell-type specific genes are organized into the same or different speckles and if the organization shows any cell type specificity. Nucleoli are involved in transcribing ribosome DNAs and ribosome assembly. Although nucleoli are mostly found away from the nuclear periphery, the nucleolus-associated chromatin domains (NADs) largely overlap with LADs. Additionally, lamins, the major structural component of the nuclear lamina, have been found to be associated with nucleolus (4, 5), but it is unclear whether this can fully explain the overlap observed between NADs and LADs. Moreover, how NADs and LADs are organized with respect to one another and to the rest of the genome is poorly understood.

Many efforts have been devoted to understanding nuclear and genome organization. Among the numerous methods developed, the imaging approaches, such as fluorescence in situ hybridization (FISH) to visualize DNA, RNA, and immunostaining to visualize proteins, allow simultaneous visualization of proteins, DNA, and RNA transcripts in individual cells. Whereas imaging approaches are important to understand the heterogeneity of nuclear and genome organization and transcriptional activity in cells, they are limited to regions of the nucleus where the probes are designed. Recent advances in barcoding and iterative hybridization approaches have enabled imaging of thousands of genomic loci in the same cell (6, 7), but the resolution is still limited.

To overcome the limitations of imaging approaches, direct DNA sequencing has been used to study genomic regions associated with specific nuclear substructure, transcription factors, or epigenetic modifications. The Chromatin ImmunoPrecipitation followed by massive parallel sequencing (ChIP-seq) (8) has been used extensively to understand epigenetic modification and transcription factor interaction with chromatin, but the method does not offer information on 3D chromatin organization or sub-nuclear structures. Several methods, including DNA adenine methylase identification (DamID) (1), nucleolar sequencing (2), Ascobate Peroxidase (APEX)-based map of LADs (9), and Tyramide-Signal Amplification (TSA)-seq (3), have been developed to map the genomic regions associated with nuclear lamina, nucleolar, and nuclear speckles. Finally, chromatin conformation capture based methods, including Hi-C (10), were developed to identify interactions between different chromatin regions, which allowed the reconstruction of the 3D chromatin interaction and folding. However, more computational efforts are still needed to fill in the gaps between DNA-DNA interaction profile and nuclear substructure-associated chromatin domains.

The A/B compartment concept was introduced in the first Hi-C study. By applying principal component analysis to the distance-normalized Hi-C interaction profile, Lieberman-Aiden et al discovered that the genome can be separated into the two large compartments, namely the A- and B-compartment (10). Genomic regions in the A-compartment had a higher chance to interact within the A-compartment, while genomic regions in the B-compartment had a higher chance of interactions with one another. By analyzing epigenome, transcriptome, and LADs in A/B compartments, it became clear that B compartments contain LADs and additional heterochromatin regions, while the A compartments were enriched for active euchromatin. We developed a model-based method, CscoreTool, to provide A-B compartment inference at higher resolution, better accuracy and lower computational cost (11, 12). These compartment studies suggest that the 3D chromatin interactions mapped by Hi-C can be used to define local chromatin compartments that share high interactions within each compartment.

To reveal more detailed 3D chromatin compartments, Rao et al used high-resolution Hi-C and Gaussian Hidden Markov Model-based clustering method to generate 6 sub-compartments and their further analyses linked some of these sub-compartments to the LADs, NADs, and speckle-associated chromatin (13). Although the algorithm used by Rao et al represents an important progression in identifying the sub-compartments of the genome, there are a number of limitations. For example, the method arbitrarily separates the genome into “even” and “odd” chromosomes and clustes them separately. The compartments inferred from the even and odd chromosomes may not match each other. Additionally, the method does not take into consideration the heterogeneity of genome interactions in individual cells known to exist even within a given cell type. Another recent method, called SNIPER (14), used a neural network to “impute” low-depth dataset and inferred sub-compartments. However, this method is supervised and depends on Rao et al’s results using the GM12878 dataset. Thus, SNIPER suffers the same limitations, and it could not be used to infer more than 6 sub-compartments or be applied to other organisms without a training dataset.

Here we introduce CscoreTool-M, a model-based tool to infer multiple sub-compartments of the genome. CscoreTool-M extends the model used in CscoreTool to accommodate more sub-compartments. The compartment scores (Cscores) provided by this method are equal to the probability (or percentage) of cells within the population that a genomic region is located in certain sub-compartments. We show that CscoreTool-M can reveal the heterogeneity of genomic sub-compartment relations within the cell population, and it offers a great opportunity to define finer sub-compartment organizations of the genome that are directly related to nuclear sub structures and their functions.

## Methods

### Modeling sub-compartments based on Hi-C dataset obtained from cell populations

#### 1. Modeling sub-compartments in population Hi-C data

We assume that there are totally *M* sub-compartments in the nucleus, and the cell population can be heterogeneous, with each genomic region *i*located to sub-compartment *k* at a probability *P*_*ik*_,

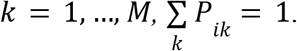

Similar to Rao et al, Our model is based on *trans*-interactions (inter-chromosome interactions). Compared to *cis*-interactions (intra-chromosome interactions), *trans*-interactions are not affected by topological domains or other local constraints. Thus, the interaction frequencies are mainly determined by compartment structure.

The model is based on three hypotheses:

Hypothesis 1: For two different genomic loci, their nuclear sub-compartment locations in cells are independent.

Hypothesis 2: Two genomic loci located at different nuclear sub-compartments in one cell do not have 3D interaction in that cell.

Hypothesis 3: Two genomic loci located at the same sub-compartment in one cell have a constant probability of interaction.

Since we only use *trans*-interactions in the analysis, local structures such as TADs would not violate our hypotheses. However, one potential violation of Hypothesis 1 is the Rabl effect in the nucleus. The Rabl effect causes centromere-centromere or telomere-telomere interactions among different chromosomes. These interactions would violate Hypothesis 1 and make the *trans*-interactions at the centromeres or telomeres dependent. To account for the Rabl effect, we add a modification in our model. Assume the Rabl effect can be modeled by a term *R*_*ij*_, then based on hypothesis 1, the probability that two genomic loci are in the same compartment in a cell is:

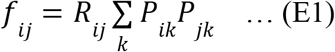

Besides the Rabl effect, these three hypotheses may still be violated under various other conditions, and we will discuss the conditions when they are violated and the consequences in section 4 below.

Based on hypothesis 2 and 3, the total number of interactions between two genomic loci should be in proportion to the probability that they are in the same genomic sub-compartment. Hi-C experiments are further affected by chromatin accessibility, genome mappability, ligation and PCR efficiency. All these complication factors are summarized as a bias factor *B*_*i*_ for each genomic locus (15), and the total interacting fragments in the Hi-C library could be modeled as:

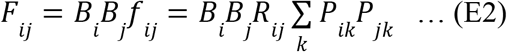

We further let *C*_*ik*_ =*B*_*i*_*P*_*ik*_, and get

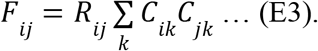

The Hi-C interaction reads between genomic loci *i* and *j, N*_*ij*_, can be modeled as a Poisson distribution with parameter *F*_*ij*_, so we get the final likelihood function:

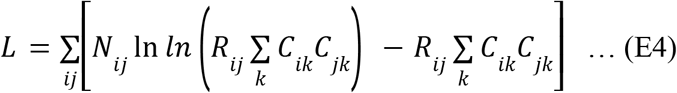

As described above the Rabl effect is simulated by a simple two-parameter model:

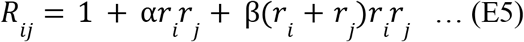

Here *r*_*i*_ corresponds to the relative location of genomic locus *i* on the chromosome. It ranges from −0.5 (centromere) to 0.5 (telomere). α is the second-order coefficient modeling how strong the Rabl effect is, while β is the third-order coefficient modeling the skew (whether the centromere or telomere side has stronger interaction).

We can infer α, β, and *C*_*ik*_ and by maximizing the likelihood function in E4 with constraints

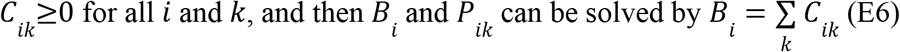

and

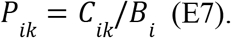

If the Rabl modifier *R*_*ij*_ is ignored, maximization of the likelihood function (E4) is in its form equivalent to symmetric non-negative matrix decomposition of the count matrix *N*≈*CC*^*T*^ with Kullback-Liebler divergence. However, as we only use trans-interactions in the analysis, only inter-chromosome genomic loci pairs are considered in calculating the divergence. Thus, this is not a standard non-negative matrix factorization problem, and we design a specific optimization algorithm to solve it as below.

#### 2. Optimization algorithm

The original optimization problem is non-convex and difficult to solve. But note that when α, β, and all the other *C*_*j*_ are all fixed, the part of *L* that is relevant to *C*_*i*_ can be written as

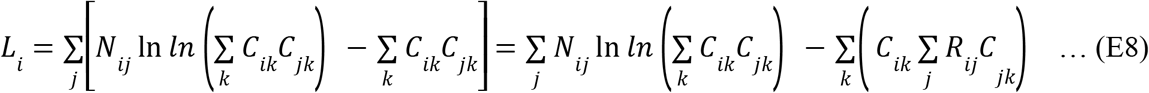

We can get the gradient and Hessian:

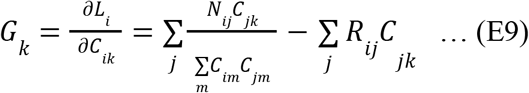

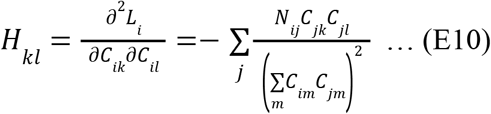

Note that − *H* is a positive-definite matrix, which means that − *L*_*i*_ is a convex function of *C*_*i*_. Since the constraints *C*_*ik*_ ≥0 are also convex, this is a standard convex optimization problem that can be solved by standard algorithms such as the interior point method. Specifically, we used the primal-dual interior point method with Newton search (16) to solve it. By fixing all the *C*_*j*_ on other chromosomes, all the *C*_*i*_ on one chromosome can be optimized in parallel.

To optimize α and β, we have the gradient

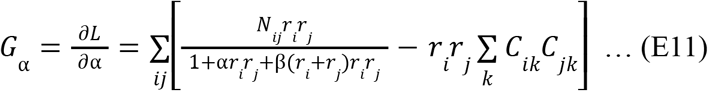

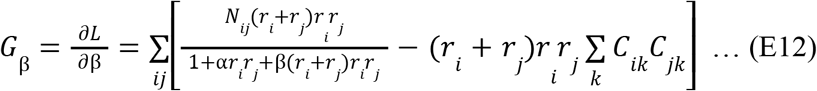

And the Hessian matrix:

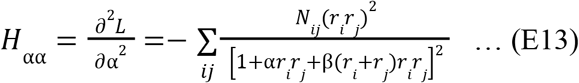

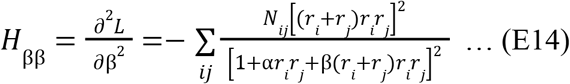

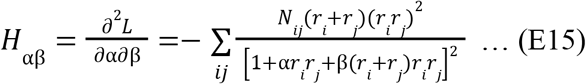

Since − *H* is also a positive-definite matrix, − *L* is also a convex function for the vector (α, β) when all the *C*_*i*_ are fixed, and the optimization problem can be solved by Newton method.

Finally, the whole algorithm works by sequentially optimize *C*_*i*_ on each chromosome, and then optimize α, β. By iterating this, each time the likelihood function will only increase so the algorithm will finally converge to a local maximum. But since the likelihood function is not overall convex, a global maximum is not guaranteed. When the number of sub-compartments *M* is large, the likelihood landscape is quite complicated, and the algorithm often converged to a local maximum. In this case, we use multiple random starting points and compare the final likelihood value to select the best one.

#### 3. Excluded regions and region groups

The human genome assembly is not perfect, and the real genome may be different among cell lines. Some of the regions have been identified to have abnormal results in genome sequencing experiments and are “blacklisted” (17). We also excluded these blacklisted regions from our analysis. All blacklisted regions will have all their ‘*C*_*i*_’s set to 0.

Moreover, there are genome translocations which make *trans*-interactions *cis*, and resulted in much more interactions than expected because genomic regions in the same chromosome interact at much higher frequency. These regions will form a separate compartment in the result because of the high interaction frequency and are easy to identify from normal compartments. When such a translocation compartment is identified, a “blacklisted region group” can be made, and in the algorithm, any pairs of regions within a blacklisted group would not be used in the analysis.

Which means in the equation E4, only those region pairs not within the same blacklisted group will be used.

#### 4. Additional conditions when the model hypotheses can be violated and the consequences

We have introduced the three hypotheses for the model above. We outline them here and discuss them one by one.

Hypothesis 1: For two different genomic loci *i* and *j*, their nuclear sub-compartment locations in individual cells are independent.

It is easy to imagine that if the two genomic loci are close to each other, like in the same topological-associated domains, the hypothesis will not be satisfied. Therefore, this hypothesis will not hold for all pairs of genomic loci. However, if we only consider trans-interactions, the locations of two genomic loci on different chromosomes should be mostly independent.

One possibility we do need to consider is that if the cell population is structurally heterogeneous, that is, it is a mixture of two or more different cell populations, which could be either different cell types or different cell states, then the independency hypothesis may be violated.

Consider there are two different cell populations, then equation E3 becomes

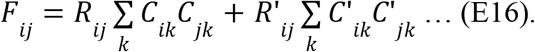

In this case, the equation E3 still largely maintains its form if the Rabl effects are similar. Only the interpretation of the *C*_*i*_values need to be changed, which means E6 and E7 may not hold. This means we will need a more careful interpretation of the C-scores when dealing with a mixed cell population. On the positive side, this means our method can identify structural heterogeneity in the cell population.

Hypothesis 2: Genomic loci located at different nuclear sub-compartments in one cell do not have 3D interaction in that cell.

In the model, the nuclear sub-compartments are corresponding to physical locations within the nucleus. Therefore, the frequency of interactions between genomic loci in different sub-compartments should be very low. However, when the two loci are located at the boundary of two neighboring sub-compartments, they should have a low chance of interacting with each other. However, when the interaction frequency is low compared to intra-compartment interactions, this will cause only minor effects.

Hypothesis 3: Genomic loci located at the same sub-compartment in one cell have a constant probability of interaction.

This hypothesis is the most likely to be violated since it is very likely DNA sequences in different nuclear sub-compartments will have very different organization and structures, and thus different physical properties and within-compartment interaction frequencies. For example, highly active genomic regions likely have higher trans-interaction frequency than condensed heterochromatin. Therefore, our hypothesis has a limitation of over-simplification, and we need to consider the consequences when this hypothesis is violated. Assume for each sub-compartment *k*, the interaction frequency has a modifier *S*_*k*_, then we have.

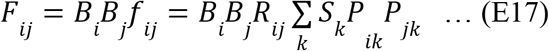

Let 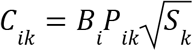 then we still have 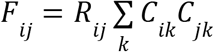 This means the equation E3 still holds in its original form, only the interpretation of *C*_*ik*_ changes. When the interaction is stronger in one sub-compartment, it will cause globally higher *C*_*ik*_ values in that sub-compartment, while weaker interactions will cause lower *C*_*ik*_ values. This will skew the prediction of *P*_*ik*_ to globally higher/lower probability. The relative higher/lower probability of subcompartment localizations between regions should still hold.

#### 5. Simulated Hi-C data generation

As discussed above, the likelihood function (E4) is complicated and non-convex. Though we can try multiple random starting points, it is uncertain whether the algorithm is robust enough to converge close to the correct optimum values or other sub-optimal local minima. To test the robustness of the method, we performed the following simulation tests.

The first simulation was performed according to our generative model with given model parameters. For each inter-chromosomal pair of genomic regions *i* and *j*, the Rabl effect *R*_*ij*_ is first calculated using equation E5. Then the expected number of interactions between the two regions can be calculated with equation E2. Finally, the number of observed interactions is generated with Poisson distribution using the expected number of interactions as the parameter.

If interactions exist, then the interactions will be output into the simulated Hi-C dataset. After iterating inter-chromosomal pairs, a full Hi-C dataset is simulated. Then CscoreTool-M is run on the simulated Hi-C data to infer compartment scores.

For low-depth dataset simulation, a full Hi-C dataset is first simulated, then a low-depth dataset is generated by down-sampling the full dataset to the lower depth.

For mixture dataset simulation, two Hi-C datasets are first simulated independently according to the two sets of parameters. Then the high-depth dataset between the two was down-sampled to match the depth of the lower one. Finally, the two Hi-C datasets were merged to become one mixed dataset.

## Results

### Validation on simulated data

To test whether CscoreTool-M can infer the correct sub-compartments, we first performed a simulation test using the GM12878 cell Hi-C data (13) as a template. The purpose of this simulation is to test whether the optimization algorithm can correctly estimate the original parameters when all the hypotheses fit. We first ran CscoreTool-M on the GM12878 cell Hi-C data to estimate compartment scores for 5 sub-compartments at 100kb resolution using the *trans* interactions (Mod1 to Mod5 on Figure 1A). Then we simulated Hi-C *trans* interactions at the same depth as the original dataset (1x dataset) using the GM12878 compartment scores as model parameters. We next applied our algorithm on the simulated dataset to infer compartment scores (Sim1 to Sim5 on Figure 1A).

**Figure 1.**
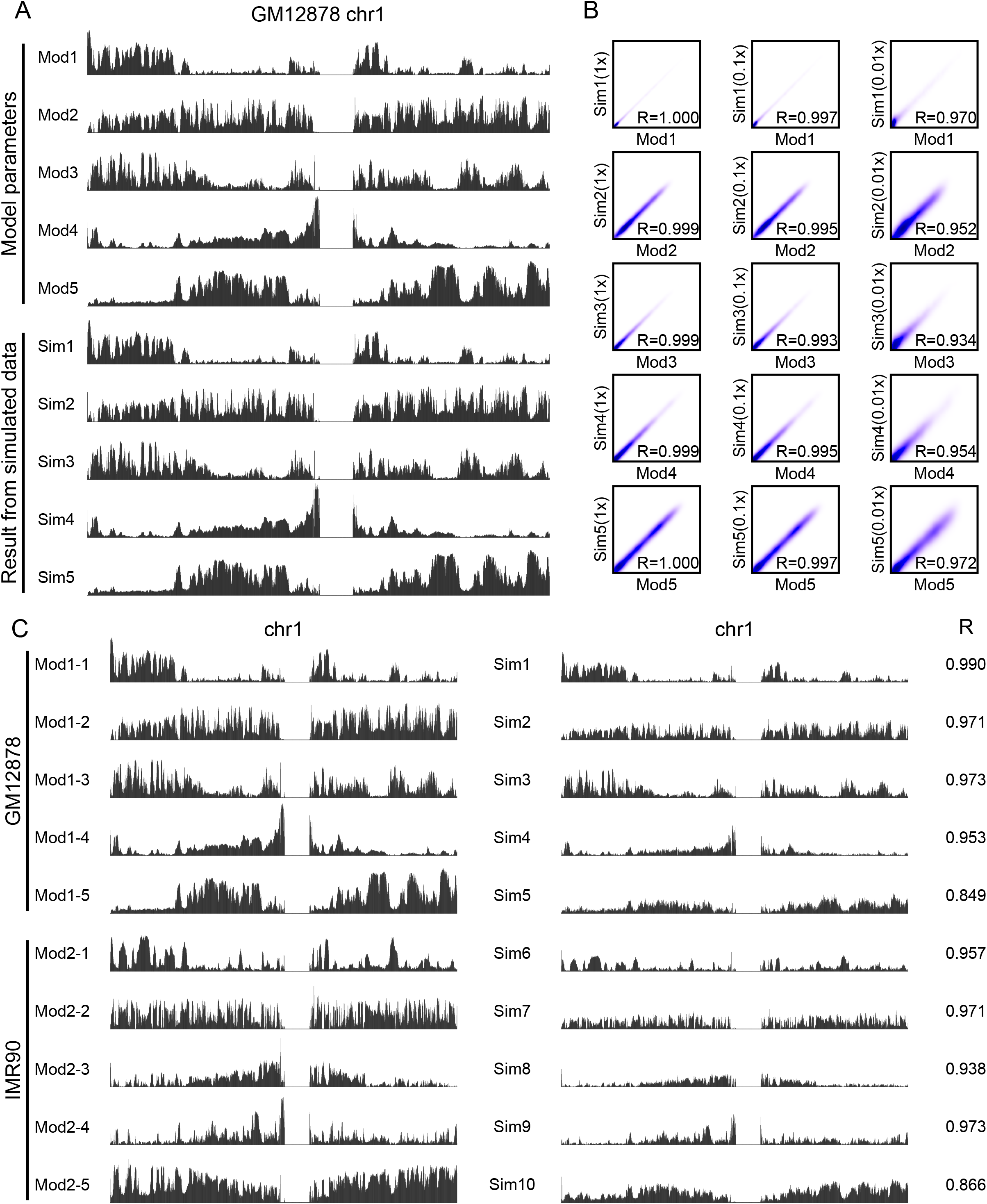
Simulation test using GM12878 cell and IMR90 cell Hi-C datasets. A. Genome browser view of compartment scores on simulated data (sim1-sim5) compared to the model values (mod1-mod5). Compartment scores on Y axes range from 0 to 1. B. Contour plots showing the compartment scores calculated on full (1x), low (0.1x), and ultralow (0.01x) depth simulated datasets compared to the model values. R: Pearson correlation coefficient. All X and Y axes compartment score values range from 0 to 1. C. Genome browser view of compartment scores on simulated mixed dataset compared to the model values. R: Pearson correlation coefficient. All Y axes range from 0 to 1.

The chromosomal view of the inferred compartment scores shows high consistency with the model parameters (Figure 1A). To quantify the divergence between the inferred compartment scores and the model parameters, we calculated the correlation coefficient at each genomic locus. High correlation is observed across all chromosomes (Figure 1B).

To further test the performance of our method at lower read depth, we generated 0.1x and 0.01x datasets by randomly selecting 10% and 1% from the original dataset, respectively, and calculated the correlation coefficients. The correlation is weaker on low-depth datasets (Figure 1B), but it is still higher than 0.9 even on the 0.01x dataset.

The theoretical analysis (see Methods section) suggested that CscoreTool-M can be used to deconvolve mixed cell population. To test this, we simulated two Hi-C datasets using model parameters for GM12878 (Mod1-1 to Mod1-5) and IMR90 Hi-C data (Mod2-1 to Mod2-5) (Figure 1C). We then mixed these two datasets and applied our algorithm to infer 10 sub-compartments (Sim1 to Sim10 on Figure 1C). We found 5 sub-compartments (Sim1 to Sim5) resembled the GM12878 model well (Mod1-1 to Mod1-5), while the other 5 compartments (Sim6 to Sim10) have a similar profile to the IMR90 model (Mod2-1 to Mod2-5). This result shows that our algorithm can infer cell-type specific sub-compartments in mixed cell types.

### Comparisons with the published multi-compartment analyses

To further test the performance of our CscoreTool-M, we compared our results with the published multi-compartments inferred by Rao and colleagues (13). Rao et al inferred 6 sub-compartments as A1-A2 and B1-B4 in GM12878 cells. A1 and A2 are both enriched for active chromatin features such as H3K4me3, H3K27Ac, H3K36me3 and DNAse I hypersensitivity. A1 has higher enrichment of active markers than A2 and A1 overlaps with nuclear speckle domains (3). B1 to B4 contain all repressed chromatin domains and are depleted of active epigenetic markers. B1 is enriched with H3K27me3 which is regulated by the Polycomb Repressive Complexes. B2 and B3 are both enriched for LADs (determined by the lamin-A/C ChIP-seq), with B2 enriched NADs (13) while B3 is not. This suggests that B2 represents NADs while B3 represents LADs. B4 only covers a very small proportion of the genome on chr19. Thus, we excluded B4 and only compared our 5 sub-compartments inferred by CscoreTool-M with the five compartments inferred by Rao et al.

We use the same notation as in Rao et al (RaoA1, RaoA2, RaoB1, RaoB2, and RaoB3) for our 5 sub-compartments but add “C5” before the compartment names (C5A1, C5A2, C5B1, C5B2, and C5B3) to indicate our CscoreTool-M modeled 5 sub-compartments. Figure 2A-B shows our 5 sub-compartment plots based on CscoreTool-M and color-coded bar-tracks for the compartments assigned by Rao et al. In general, our compartment scores show peaks at Rao’s assigned compartments, indicating that our compartment inference is largely consistent with the compartments indicated by Rao. It is noteworthy that there are also regions showing inconsistencies. Figure 2A and B show two regions on chr10 and chr12 have C5B1 sub-compartment scores peaks, while Rao et al. predicted these regions to be in A1. To further investigate which sub-compartments these regions should be in, we looked at gene expression and epigenetics features along these regions. Our C5B1 peaks show a depletion of H3K27Ac and reduced gene expression, but enriched for H3K27me3, which are typical B1 features instead of A1.

**Figure 2.**
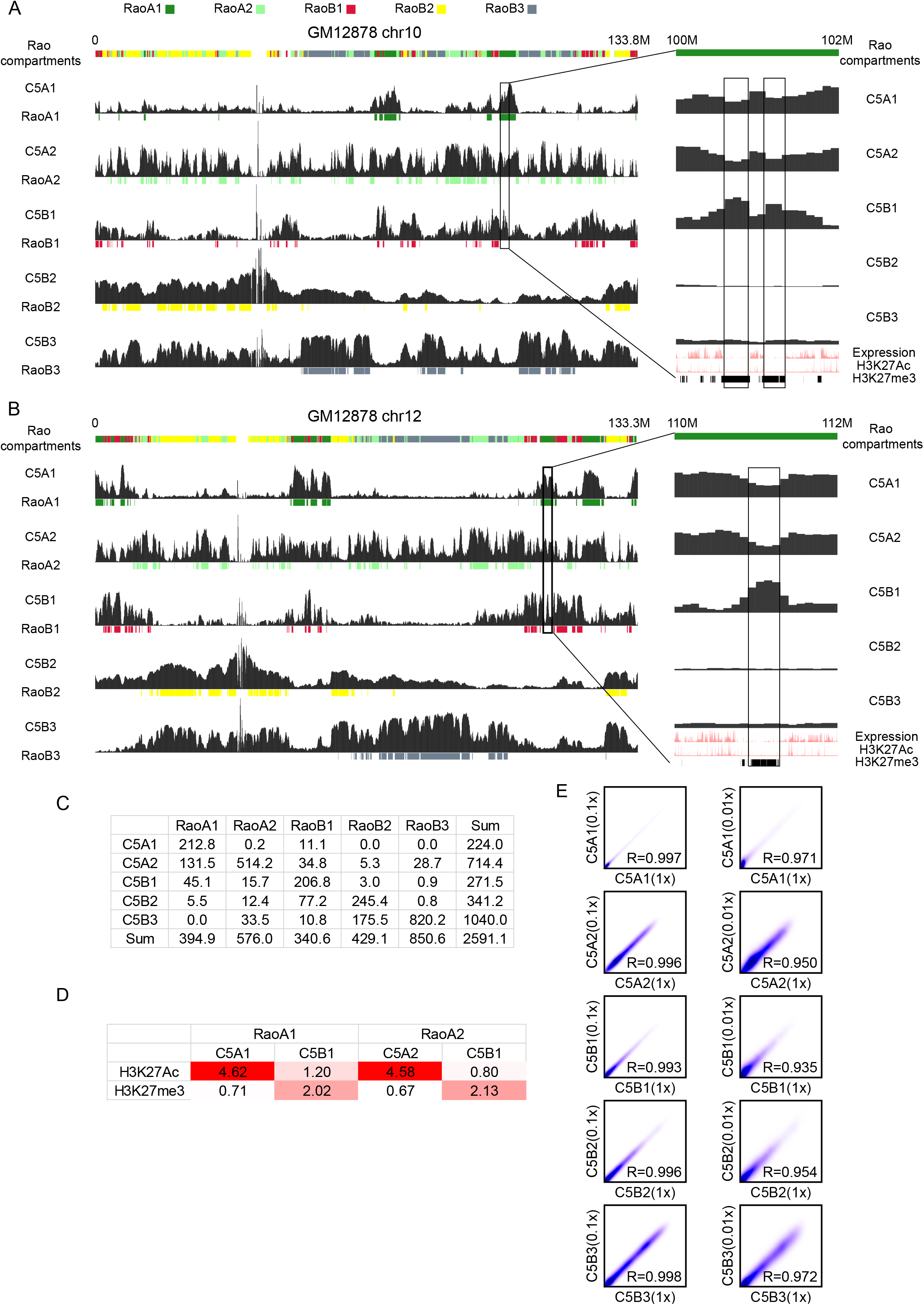
Comparisons of compartments modeled by CscoreTool-M against Rao’s sub compartments in GM12878 cells. A-B. Genome browser view of chromosome 10 (A) and 12 (B) showing the compartment scores tracks by CscoreTool-M and compartments (indicated by different colored blocks) by Rao et al. Example regions in black rectangles showing discrepancies between CscoreTool-M and Rao et al are expanded and shown at the right panels. All Y axes plot for compartment scores range from 0 to 1. C. Comparisons between compartments inferred by CscoreTool-M and Rao et al. Values are in million bases. D. Enrichment of H3K27Ac and H3K27me3 peaks in regions agreed or disagreed inferred by CscoreTool-M or Rao et al. H3K27Ac and HeK27me3 data are from UCSC genome browser ENCODE tracks. E. Contour plots showing the compartment scores calculated on low (0.1x) and ultralow (0.01x) depth datasets compared to the original dataset. All compartment score values on X and Y axes range from 0 to 1.

To further analyze whether the difference in A and B sub-compartment assignment by Rao and by our CscoreTool-M also happens on other chromosomes, we assigned the 5 compartments based on the highest compartment score at each genomic region across the genome and compared these to the compartments assigned by Rao et al (Figure 2C). While the overall correspondence in A and B sub-compartments is good, there are some clear differences. For example, we found that a total of 45.1M genomic regions are assigned as A1 by Rao but are assigned as C5B1 by CscoreTool-M (Figure 2C). We further looked at the enrichment of H3K27Ac peaks and H3K27me3 peaks on these regions (Figure 2D) and found that the H3K27Ac enrichment (1.2) is just marginally higher than the global average (1.0), while much lower than the enrichment of H3K27Ac peaks on A1 regions assigned commonly by Rao and CscoreTool-M. By contrast, the H3K27me3 peaks showed much higher enrichment (2.02) than that found on A1 regions assigned commonly by Rao and CscoreTool-M (0.71), and also higher than the global average (1.0). We made a similar observation in regions assigned as A2 by Rao and C5B1 by CscoreTool-M. The C5B1 regions show lower H3K27Ac and higher H3K27me3 than the corresponding RaoA2 regions and also than global average (Figure 2D). These results show that the CscoreTool-M assigned C5B1 regions have B1 features instead of A1/A2 features. Thus, the CscoreTool-M can better predict sub-compartments in these regions.

We also tested the performance of our CscoreTool-M on low-depth Hi-C sequencing data by randomly selecting 0.1x and 0.01x of the reads from the Hi-C data for GM12878 cells to infer compartments. We then calculated the correlation coefficient on the compartments. We found all the correlation coefficients are higher than 0.9 (Figure 2E), indicating that the CscoreTool-M is robust in inferring multiple genomic compartments using low-depth Hi-C datasets.

### Inferring LAD and NAD heterogeneity within a cell population

One feature of CscoreTool-M is that it explicitly models and quantifies the heterogeneity within a cell population, which should aid the study of variations of genome organization. It is known that NADs often overlap with LADs, but the overlap only happens for some NADs. The NAD-only and the overlapping NAD/LAD chromatin regions tend to have different genomic features (18, 19). Since nucleoli are often not located at the nuclear periphery, the biological interpretation of the overlap between NADs and LADs remains unclear. The localization of lamins in nucleolus as reported by some studies (4, 5) can contribute to the NAD/LAD overlap. Additionally, only a fraction of LADs may return to the nuclear lamina after mitosis, and the ones not returning to the nuclear lamina often relocate to peri-nucleolar regions (20). In this case, the overlapping LADs and NADs regions can come from heterogeneity within the cell population, with the regions located at the nucleoli or nuclear periphery in different cells within the population.

We applied CscoreTool-M to infer LADs and NADs based on the Hi-C dataset for the GM12878 cell population. One interesting feature we noticed is that our compartment scores show clear variations among Rao et al’s B2 chromatin regions. Examples are shown for chr18 and chr19 (Figure 3B), which are known to localize preferentially to the nuclear periphery and nuclear interior, respectively (21). On chr18, about half of the chromosome was assigned as B2 by Rao et al (RaoB2), but based on Cscoretool-M these regions have similar C5B2 and C5B3 scores. Although RaoB2 and RaoB3 are suggested to represent NADs and LADs regions, respectively, our analyses indicate that the RaoB2 chromatin regions on chr18 exhibit cell-cell heterogeneity with about half of the cells having nucleolar localization and the other half localizing at the nuclear lamina. In contrast to chr18, CscoreTool-M found that the RaoB2 regions on chr19 have high C5B2 scores and low C5B3 scores, suggesting that these regions are NADs but not LADs. This correctly infers that the interiorly localized chr19 has limited LADs compared to that of chr18. Taken together, our method can infer the heterogeneity of NADs and LADs localization and quantify the different levels of heterogeneity among genomic regions within the cell population.

**Figure 3.**
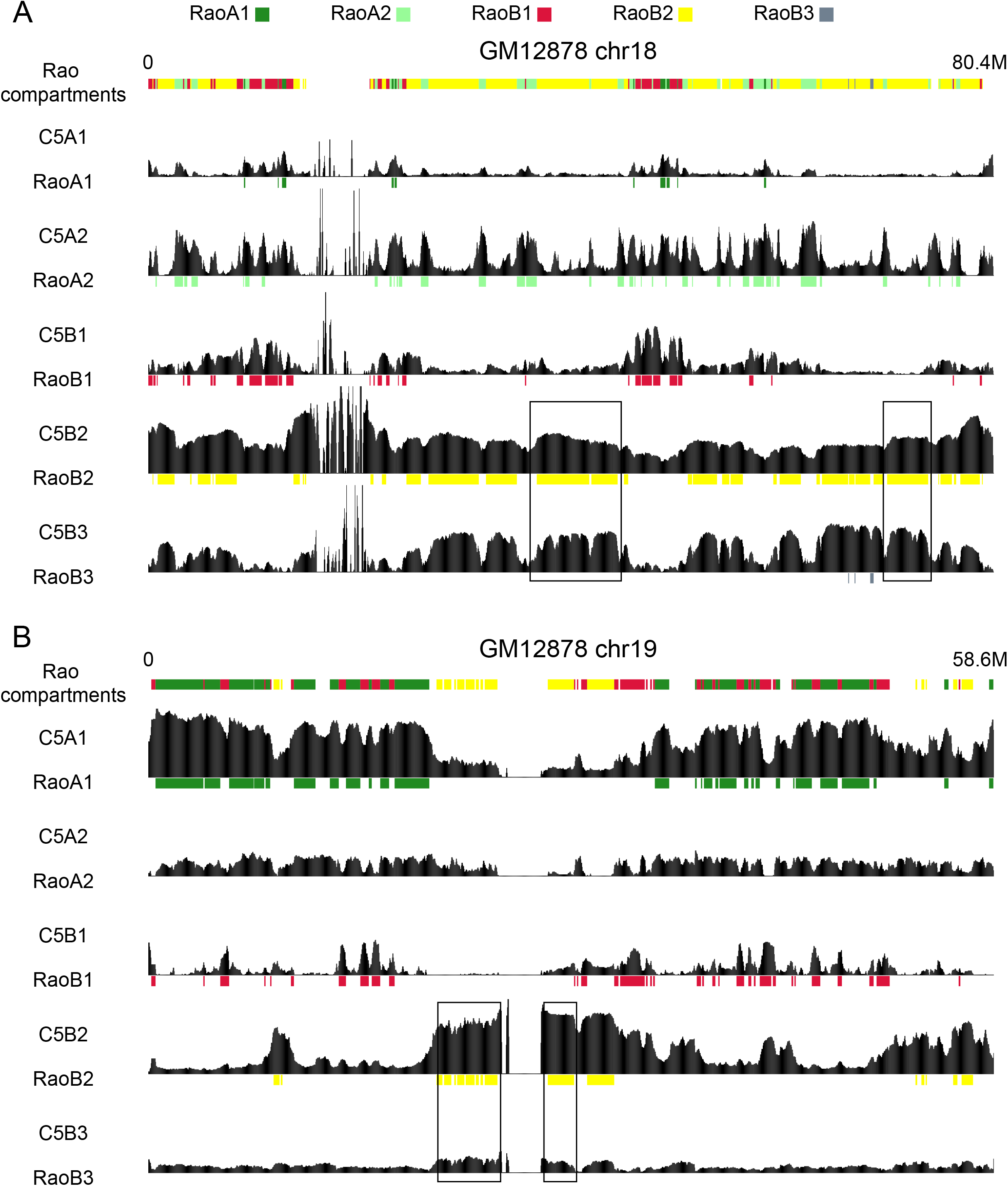
Inferring NADs and LADs in GM12878 cells. Genome browser view of chromosome 18 (A) and 19 (B) showing the CscoreTool-M compartment score tracks and the corresponding compartments by inferred by Rao et al. Black rectangles indicate two examples of RaoB2 regions showing high C5B2 and C5B3 scores on chr18 (A) or two examples of RaoB2 regions with high C5B2 and low C5B3 scores on chr19 (B). All Y axes plotting compartment scores range from 0 to 1.

### Inferring the degree of compartment heterogeneity in different cell types

Studies have revealed clear variations in 3D genome interactions in individual cells belonging to the same cell type. These variations can be purely stochastic or related to different cell states, such as different cell cycle or differentiation stages. On the other hand, a given cell type that is not proliferating and is terminally differentiated should exhibit relatively low heterogeneity in genome organization. Our CscoreTool-M should be able to infer the degree of 3D genome organizational heterogeneities among different cell types and within a population of cells belonging to the same cell type or lineage. To test this, we applied CscoreTool-M to analyze the Hi-C dataset for the isolated pure mature olfactory sensory neurons (OSN)(22). The result of the 5-compartment analyses (Figure 4A) shows sharper sub-compartment boundaries and lower background scores than those of the 5 compartments inferred based on the Hi-C datasets for GM12878 and IMR90 cells (see Figure 1C).

**Figure 4.**
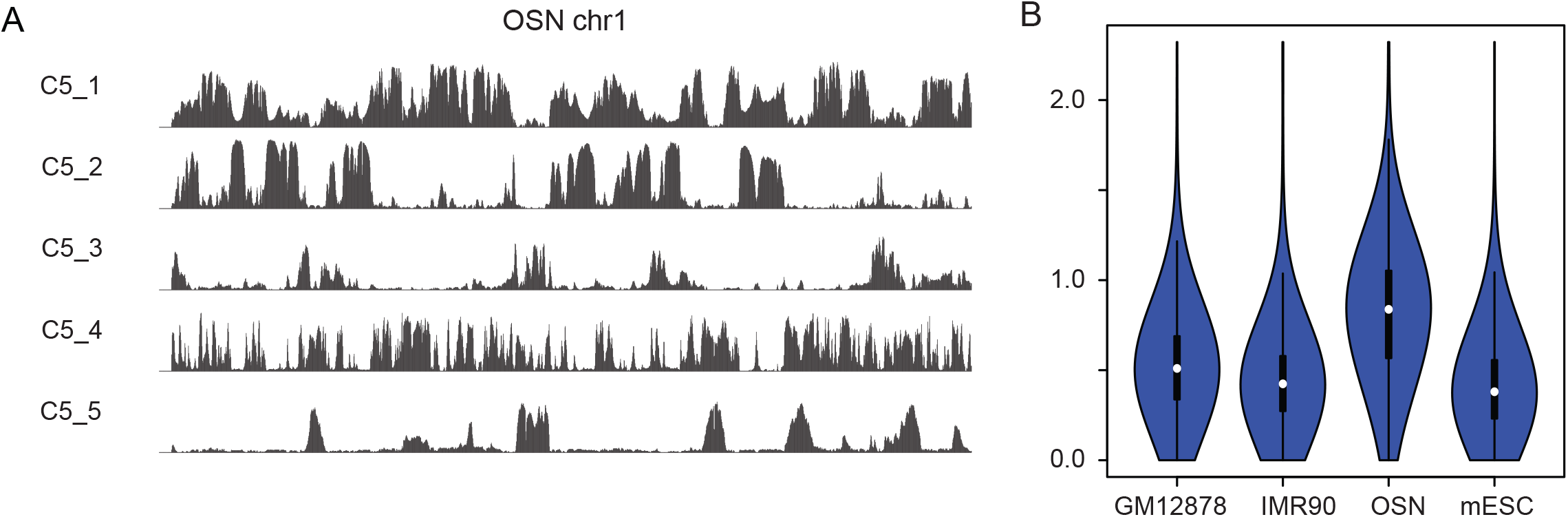
Comparisons of the level of compartment heterogeneity among different cell types. A. Genome browser view of 5 compartments for olfactory sensory neurons (OSN) on chr1 inferred by CscoreTool-M. All Y axes plotting compartment scores range from 0 to 1. B. Violin plot showing the information content calculated for each 100-kb window along the genome based on Hi-C datasets in 4 different cell types.

To further quantify the degree of heterogeneity, we calculated the information content of sub-compartment scores along the genome. The information content, defined as 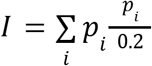 for 5 sub-compartments, is highest if the sub-compartment score is 1 for one sub-compartment and 0 for the remaining compartments (no heterogeneity), while lowest (0) if it is the same for all sub-compartments (no information). We found that the information content for the olfactory neuron Hi-C dataset is generally higher than the GM12878 and IMR90 Hi-C datasets (Figure 4B), confirming our visual inspection of the genomic tracks. Since the olfactory sensory neurons were isolated from mice, we compared the information content of the mouse olfactory neuron Hi-C with a mouse ES cell (mESC) Hi-C dataset (23). The olfactory neuron dataset still has significantly higher information content (Figure 4B). Thus, the difference is not due to the differences of cells derived from mice or humans. These results show that CscoreTool-M can be used to infer and compare genomic organizational heterogeneity in different cell types. The results also suggest that cells that continue to proliferate may result in increased genome organizational heterogeneity.

### Inferring G1 genome organization in a population of asynchronously dividing cells

In a given cell type, those cells that are proliferating can have different 3D genome interactions at different cell cycle stages, thereby contributing to the variation in chromatin organization among individual cells. Our simulation analysis suggests that the CscoreTool-M is able to deconvolve genomic organizational heterogeneity if we analyze a large number of sub-compartments. We analyzed the Hi-C dataset obtained from the asynchronously dividing HCT116 cells (24) using10 sub-compartments (Figure 5A). Recently we have developed a chromatin pull-down based Tyramide-Signal Amplification sequencing (cTSA-seq) method to analyze LADs at different cell cycle stages using the human HCT116 cell line (25). We found that during early G1 the chromatin associated with the reforming NL are the sub-telomeric regions. This observation was also observed using the novel protein A-DamID approach described by van Steensel and colleagues (26). Interestingly, one sub-compartment, HCT116 C10_7, predicted by CscoreTool-M is very similar to the early G1 LADs mapped by our cTSA-seq. The whole-genome comparison shows significant correlation between the compartment score of HCT116 C10_7 and the early G1 cTSA-seq LADs mapping (R=0.42, P<10^−16^). This result shows that the C10_7 sub-compartment can capture the LADs pattern in the early G1 HCT116 cells.

**Figure 5.**
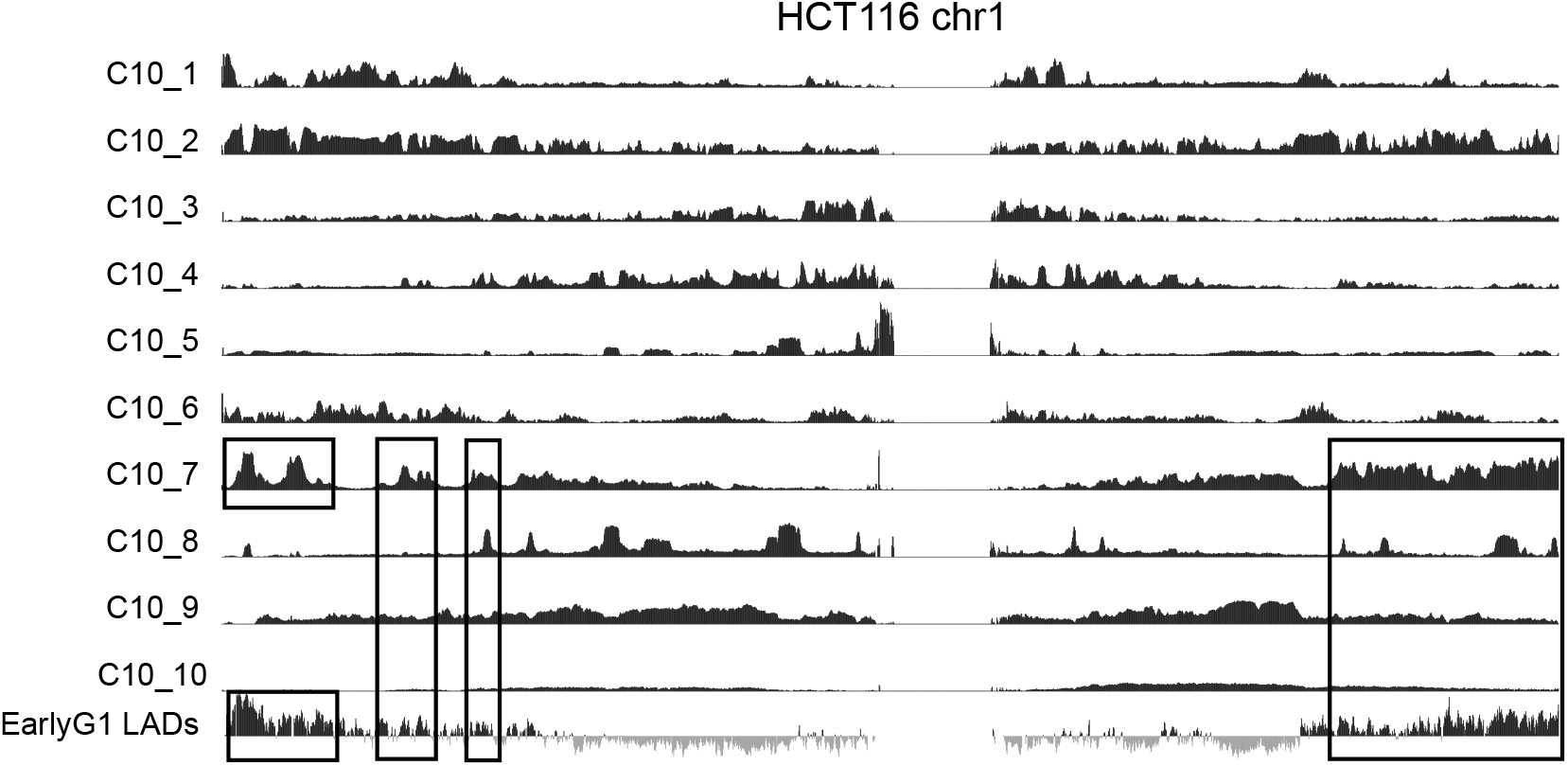
Inferring cell-cycle-related sub-compartments. Genome browser view of 10 compartments inferred by CscoreTool-M for HCT116 cells on chr1. Regions showing similarity between HCT116 C10_7 compartment and the early-G1-specific LADs determined by cTSA-seq using sorted early G1 HCT116 cells are highlighted in black boxes.

### Inferring cell-type specific functional sub-compartments

One interesting feature of the olfactory neuron is that the olfactory genes are clustered together in 3D (22). To test whether our method can find this sub-compartment of olfactory genes, we applied the CscoreTool-M on the olfactory sensory neuron Hi-C dataset to model 10 sub-compartments. We found that one of the sub-compartments, OSN C10_1, overlapped well with the olfactory genes (Figure 6A), indicating that our method can identify this functional gene cluster in one sub-compartment.

**Figure 6.**
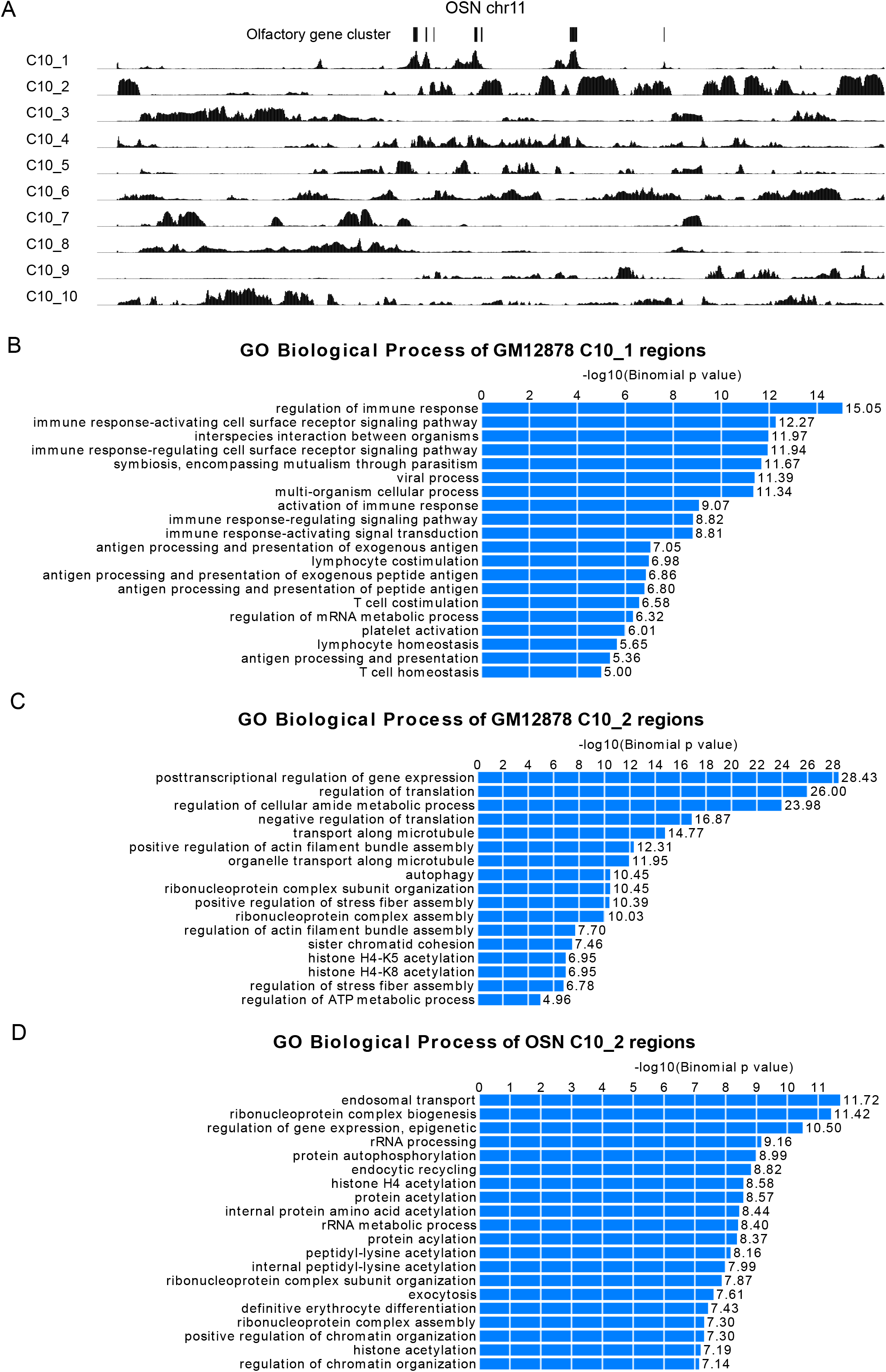
Identification of function-related sub-compartments in GM12878 and OSN by 10 sub-compartment analyses. A. Genome browser view of 10-compartments for olfactory sensory neuron cells on chr11. The olfactory gene clusters are indicated with black bars on the top. Thickness of the bars indicate the number of olfactory genes. B-C. GO-term enrichments by GREAT of genes in the compartment C10_1 (B) and C10_2 (C) compartments in GM12878 cell. D. GO-term enrichments of the C10_2 compartment in olfactory and sensory neurons (OSN).

We reason that these functional gene clusters may also exist in other cell types. We first examined the GM12878 dataset. We performed 10-compartment analysis on the GM12878 dataset and found that the GM12878 C10_1 and C10_2 sub-compartments both have good correlations to the GM12878 C5A1 compartment but are enriched for genes with different functions based on the Gene Ontology (GO) analyses. The C10_1 sub-compartment is enriched for genes related to immune function, which may be functionally important as the GM12878 cells are derived from B-lymphocytes (Figure 6B). By contrast, the C10_2 sub-compartment is enriched for housekeeping genes with functions including translation and transcription (Figure 6C). Interestingly, we found that the C10_2 sub-compartment for the olfactory sensory neurons is also enriched for housekeeping genes (Figure 6D).

Finally, we applied CscoreTool-M to infer 10 sub compartments in additional cell types with available Hi-C datasets to test if we can find 3D clustering of functionally related genes in individual genomic compartments. By analyzing mESCs, IMR90 fibroblasts, NHEK keratinocytes, and HCT116 colon cancer epithelial cells, we found a clear enrichment of housekeeping genes in two sub-compartments of mESC (Figure 7A and B), one in IMR90 (Figure 7D) and one in HCT116 cells (Figure 7E) GO analyses also revealed an enrichment of genes in the C10_3 compartment of mESC involved in embryonic development (Figure 7C) and genes in the C10_2 compartment of NHEK cells involved in keratinocyte differentiation and skin development (Figure 7F). Thus, CscoreTool-M can discover the 3D cluster of functionally related genes in different cell types.

**Figure 7.**
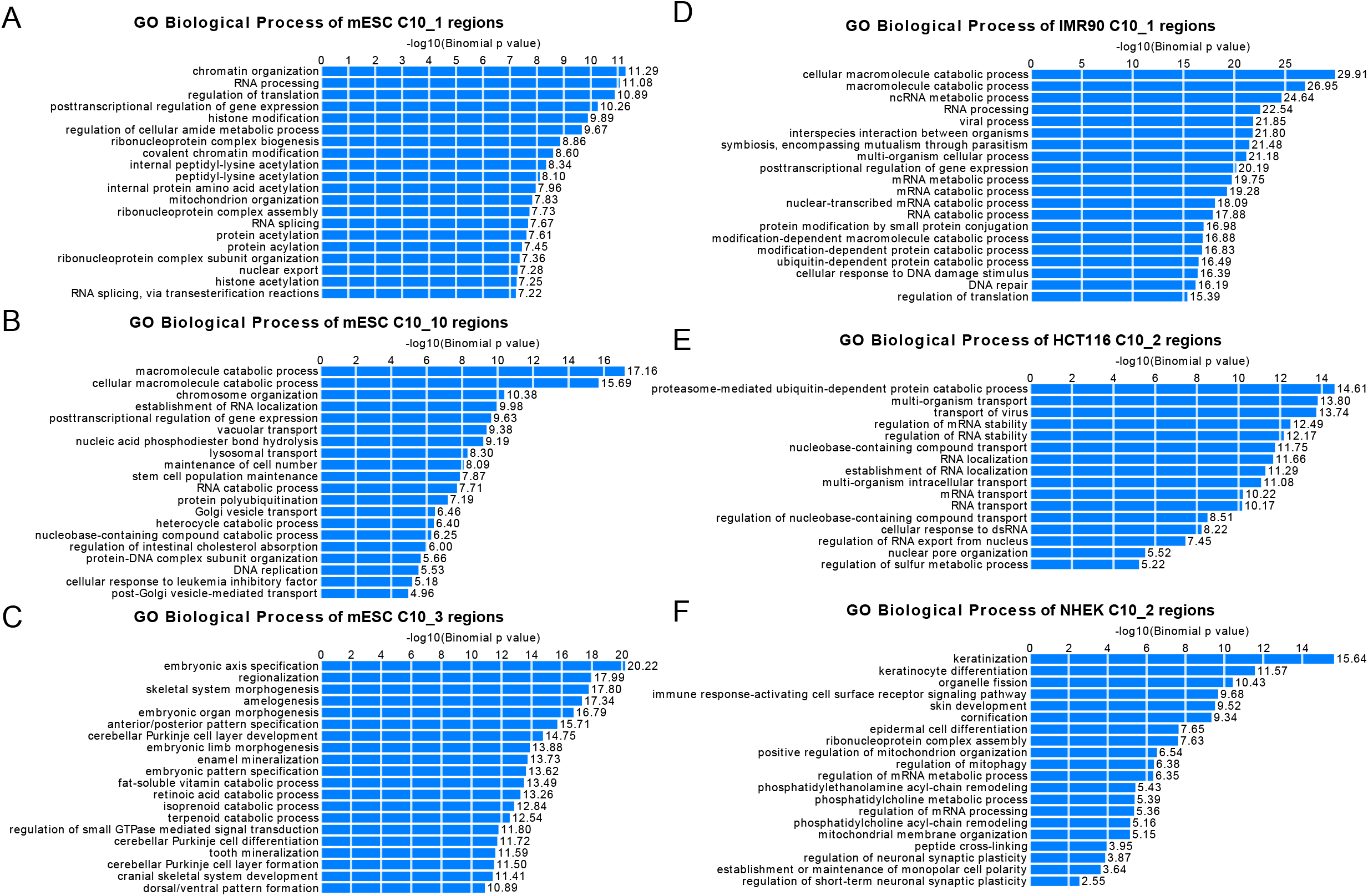
GO-term analysis of functional-related sub-compartments based on 10 sub-compartment modeling in mESCs (A-C), IMR90 (D), HCT116 (E) and NHEK(F) cells.

## Conclusions

In this study, we report CscoreTool-M, a model-based method to calculate sub-compartment scores from Hi-C data. The compartment scores calculated by Cscoretool-M for a given genomic region are directly proportional to the probability that the region is located at a specific sub-compartment. By comparing with Rao et al’s sub-compartment inference based on clustering method, we have shown that CscoreTool-M is better at differentiating sub-compartments enriched for repressive heterochromatin marked by H3K27me3 from the surrounding sub-compartments enriched for active chromatin compared to the clustering method. The compartment scores by CscoreTool-M also help quantify the levels of heterogeneity in sub-compartment localization within cell populations for different genomic regions. For example, CscoreTool-M quantifies different C5B2 and C5B3 scores among different chromosomes. By comparing between proliferating and terminally differentiated cells, we show that proliferating cells have higher genome organization heterogeneity, which is likely caused by cell-cycle stages. By analyzing 10 sub-compartments, we found a sub-compartment potentially related to early-G1 LADs in HCT116 cells, suggesting the method can deconvolve sub-compartments from asynchronously dividing cell populations. Finally, we show that sub-compartments inferred by CscoreTool-M are often enriched in housekeeping or cell-type-specific functions.

## Data Availability

The CscoreTool-M source code and sub-compartment scores analyzed in this study have been deposited at https://github.com/scoutzxb/CscoreTool-M.

## Acknowledgement

This work has been supported by the National Institutes of Health (R01GM106023 and R01GM110151 to Y.Z.).

